# In silico stress fibre content affects peak strain in cytoplasm and nucleus but not in the membrane for uniaxial substrate stretch

**DOI:** 10.1101/774976

**Authors:** Tamer Abdalrahman, Neil H. Davies, Thomas Franz

## Abstract

Existing in silico models for single cell mechanics feature limited representations of cytoskeletal structures that contribute substantially to the mechanics of a cell. We propose a micromechanical hierarchical approach to capture the mechanical contribution of actin stress fibres. For a cell-specific fibroblast geometry with membrane, cytoplasm and nucleus, the Mori-Tanaka homogenization method was employed to describe cytoplasmic inhomogeneities and constitutive contribution of actin stress fibres. The homogenization was implemented in a finite element model of the fibroblast attached to a substrate through focal adhesions. Strain in cell membrane, cytoplasm and nucleus due to uniaxial substrate stretch was assessed for different stress fibre volume fractions and different elastic modulus of the substrate. A considerable decrease of the peak strain with increasing stress fibre content was observed in cytoplasm and nucleus but not the membrane, whereas the peak strain in cytoplasm, nucleus and membrane increased for increasing elastic modulus of the substrate.

## 1. Introduction

Knowing how cells deform under different loads is crucial for improving the understanding of physiological and pathological events [1, 2]. Eukaryotic cells comprise various components (e.g. cytosol, fibrous protein networks, nucleus, cell membrane) that are dispersed heterogeneously, display non-linear behaviour [3, 4], and can be highly anisotropic [5]. The cytoskeletal components determine the mechanical properties of the cell. These properties can be quantified through experimental characterization and theoretical formulations [6, 7] and have been found to vary even for the same type of cell [8].

Various techniques have been developed to obtain the mechanical properties of cells, including micropipette aspiration [9], use of optical tweezers [10], magnetometric examination [11] and atomic force microscopy (AFM) based strategies [12].

The cytoskeleton, and particularly actin filaments, affect the morphological and mechanical properties of the cell. For example, cytoskeletal changes associated with cellular remodelling may lead to substantial changes in a cell’s mechanical properties [14–17]. Park et al. [13] examined the localized cell stiffness and its correlation with the cytoskeleton. They reported that the local variation of cytoskeletal stiffness was related to regional prestress. This study comprehensively characterized the localized variations of intracellular mechanical properties that underlie localized cellular function.

Rotsch and Radmacher [18] found a substantial reduction in the elastic modulus of fibroblasts when treated with actin disrupting chemicals, with similar findings reported by others [19, 20]. The variation in cellular stiffness has been linked to diseases [21], such as the decrease of the elastic modulus of cancerous cells by one order of magnitude compared to healthy cells [22]. With some exceptions [6, 7, 23, 24], previous studies applied homogenization to the entire cell and did not explicitly consider the effect of inhomogeneities [25]. The homogenization resulted in non-physical relationships between assessed parameters and the mechanical properties of the cell.

In computational cell mechanics, an accurate description of the cytoskeleton’s anisotropic, non-linear behaviour is desired to account for cytoplasmic inhomogeneity. Finite element methods (FEM) have been utilized to study various cell mechanics aspects [26–28]. The application of FEM to single-cell and subcellular mechanics [29–31] is still limited because of the scarcity of data on material properties and shape of sub-cellular structures. The combination of image-based geometrical modelling and FEM have recently facilitated computational models with three-dimensional (3D) cellular morphology that comprised cytoskeleton, cytoplasm, cell membrane, and nucleus [9, 10, 32–34]. However, including cytoskeletal stress fibres as discrete structural elements is one of the current challenges in computational cell mechanics. Multiscale constitutive models may offer a way to address this challenge by capturing the mechanical properties of cellular structures at the sub-cellular length scale and representing their mechanical contribution at the cellular length scale.

This study aimed to develop a multiscale computational model, combining micromechanical homogenization and finite element methods, capable of capturing the mechanical contributions of intracellular structures. The model’s feasibility was demonstrated by investigating the effects of the actin stress fibre content and substrate elastic modulus on the intracellular mechanics of a single fibroblast.

## 2. Materials and methods

### 2.1 Geometrical modelling

The geometrical model of a human dermal fibroblast developed previously [30] was utilized in the current study. In brief, human dermal fibroblasts were seeded onto fibronectin-coated sterile cover slips, stained with Alexa Fluor 568 phalloidin (Invitrogen Molecular Probes, Eugene, Oregon, USA) for actin stress fibres and counterstained with Hoechst 33342 dye (Sigma Aldrich Chemie GmbH, Steinheim, Germany) for the nucleus. Confocal images were acquired with a Zeiss 510 LSM Meta microscope at 40x magnification.

The 3D cellular geometry was reconstructed from a confocal z-stack (**Figure 1** a) with 31 images (dimensions: x = 225 μm and y = 225 μm, z = 17.4 μm, z-interval = 0.58 μm) using threshold-based segmentation of the phalloidin stain for actin fibres to represent the cytoplasm and the Hoechst stain for the nucleus (Simpleware ScanIP, Synopsys, Mountain View, CA, USA) [30]. The reconstructed cytoplasm had in-plane dimensions of 145 and 92 μm in the long and short axis, respectively, and a thickness of 14 μm. The nucleus had an in-plane diameter of 19 μm and a maximum thickness of 3 μm (**Figure 1** b). The reconstructed geometry was complemented with a membrane with a thickness of 0.01 μm [35–37] enveloping the cytoplasm (Simpleware ScanIP) (**Figure 1** c).

**Figure 1.**
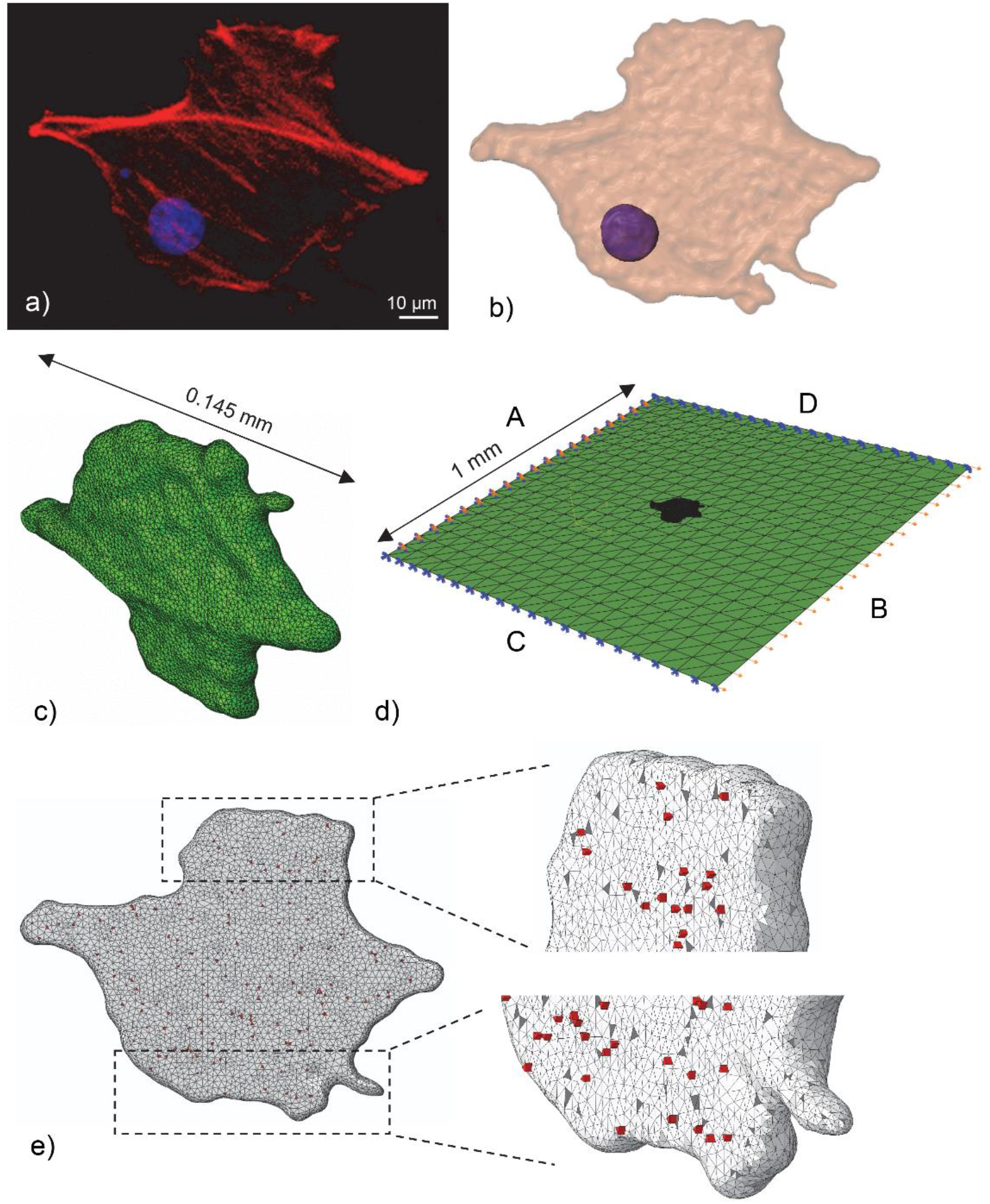
Microscopic image and computational geometry of fibroblast. a) Maximum intensity projection of confocal microscopic image stack of stained actin fibres (red) and nucleus (purple) of fibroblast (basal view). b) 3D reconstructed geometry of cytosol (transparent beige) and nucleus (purple) (apical view). c) Meshed geometry of the fibroblast and nucleus (not visible) (apical view). d) Finite element mesh of cell attached to the substrate with boundary conditions of the substrate: Edge (A) fixed in all directions; uniaxial quasi-static displacement applied in the normal direction to opposite edge B; edges C and D free in the displacement direction and fixed in the normal direction. e) Illustration of focal adhesion cohesive elements (red) in two regions of basal cell surface. The substrate is not shown for clarity. The nodes of the cohesive elements did not conform with nodes of the cell and substrate meshes. Tie constraints were used to define contact between focal adhesions and cell and substrate, respectively.

### 2.2 Finite element modelling

#### 2.2.1 Mesh generation

The cellular components were meshed with shell (membrane) and tetrahedral elements (cytosol and nucleus), see **Table 1**, converted to volumes and defined as separate element sets. No-slip conditions were enforced at interfaces between the cellular components in Simpleware ScanIP. The meshed cell geometry was imported into Abaqus CAE (Dassault Systèmes, RI, USA) and assembled with a 1 x 1 mm flat substrate to simulate exposure of the cell to substrate stretch [28], see **Figure 1**(d).

**Table 1.** Types and number of elements for each component in the cell-substrate finite element model

| Component | Type of element* | Number of elements |
| --- | --- | --- |
| Nucleus | C3D4 | 1075 |
| Cytosol | C3D4 | 47029 |
| Plasma membrane | S3R | 14832 |
| Elastic substrate | S3 | 800 |
| Focal adhesion | COH3D6 | 248 |

\*C3D4 - four-node tetrahedral element, S3 - triangular three-node shell element, S3R - triangular three-node triangular shell element with reduced integration, COH3D6 - six-node three-dimensional cohesive element

For the cell-substrate attachment, 124 focal adhesions (FA) with a thickness of 1 μm [38–41] and average contact area of 1 μm^2^ [41] were randomly distributed across the basal cell surface [30] and meshed with cohesive elements [28, 42–44] (**Figure 1** e). The number and types of elements of the various parts of the model are provided in **Table 1**.

#### 2.2.2 Contacts, boundary conditions and loading

Tie constraints were used between the cohesive elements and the cell membrane and elastic substrate, respectively. The cell was positioned in the substrate centre with sufficient distance to the substrate boundaries to neglect edges effects (**Figure 1** d). One edge (A) of the substrate was fixed in all directions, and a uniaxial quasi-static displacement was applied normally to the opposite edge (B). The other two edges (C and D) remained free in the displacement direction and fixed in the normal direction. The applied displacement generated a uniform deformation field in the substrate with a tensile strain up to 10% (i.e. stretch λ = 1.1).

#### 2.2.3 Material Properties

The finite element simulations were limited to strain field. The cell membrane was represented as isotropic linear-elastic, whereas nucleus and cytosol were assumed to be isotropic hyper-elastic compressible and described with a Neo-Hookean strain energy function [45–47]:

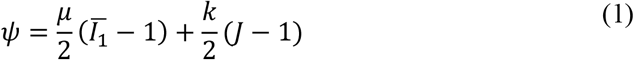

where *ψ* is the strain energy per volume reference unit, μ is the shear modulus, and k is the bulk modulus. 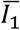 and J are the first and third invariant of the left Cauchy-Green deformation tensor, **P**, given by:

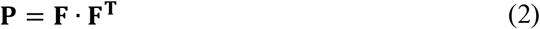

where **F** is the deformation gradient tensor [48]. The substrate and focal adhesions were represented with an isotropic linear-elastic material model. All materials parameters are provided in **Table 2**.

**Table 2.** Mechanical properties of cell components, substrate, and focal adhesions.

| Component | Constitutive law | Parameter |
| --- | --- | --- |
| Nucleus | Hyperelastic | $\mu = 1.7$ kPa, $k = 16.4$ kPa [49] |
| Cytosol | Hyperelastic | $\mu = 1$ kPa, $k = 9.7$ kPa [45] |
| Stress fibre | Linear-elastic | $E = 1.45$ MPa, $\nu = 0.3$ [32] |
| Membrane | Linear-elastic | $E = 7$ kPa, $\nu = 0.45$ [50] |
| Focal adhesion | Linear-elastic | $E = 10$ kPa, $\nu = 0.3$ [51, 52] |
| Elastic substrate | Linear-elastic | $E_{\text{sub}} = 0.01, 0.14, 1, 10$ MPa, $\nu = 0.45$ |

### 2.3 Micromechanical homogenization of cytoplasm

The components of the cytoskeleton include actin filaments, intermediate filaments, and microtubules. The cell can reorganize the cytoskeletal filaments according to its microenvironment, thus changing its mechanical properties. For simplification, intermediate filaments and microtubules were disregarded in the current study involving small cell and substrate tensile deformations. This omission was deemed acceptable since (i) the intermediate filaments bear tension loads predominantly for large cell deformations and only to a small extent for small deformation [53, 54], and (ii) microtubules provide structural support primarily for compressive loads [55]. Stress fibres are contractile bundles of actin filaments [56] with a diameter in the range of tenths of microns. Experimental and theoretical studies have shown that cellular behaviour, such as migration, depends on the organization of actin stress fibres [57, 58]. in this study we consider the stress fibres as randomly distributed in the cytoplasm, satisfying the continuum hypothesis.

The micromechanical homogenization allows obtaining the cytoplasm’s effective mechanical properties by treating the cytoplasm as a composite of cytosol and stress fibres (see **Figure 2**). Homogenization is achieved by substituting the heterogenous microstructure with an equivalent homogenized structure. The micromechanical homogenization of the material properties is obtained considering a representative volume element (RVE). The RVE is a statistical representation of the material’s microstructure and needs to provide sufficient details of the micro-fields to ensure accurate sampling of the entire domain. The micro-length scale of the structure is represented by the RVE, which is small (*l* < L) compared to the macrostructure (i.e. the cell) and assumed to approximate the mechanical properties of the macrostructure. The RVE is, however, very large (*l* >> *ℓ*) compared to the microstructural elements (i.e. the stress fibres) and not suited capture the microstructure in sufficient detail. The cytoplasm’s effective mechanical properties are then obtained based on the volume fraction of the stress fibres.

**Figure 2.**
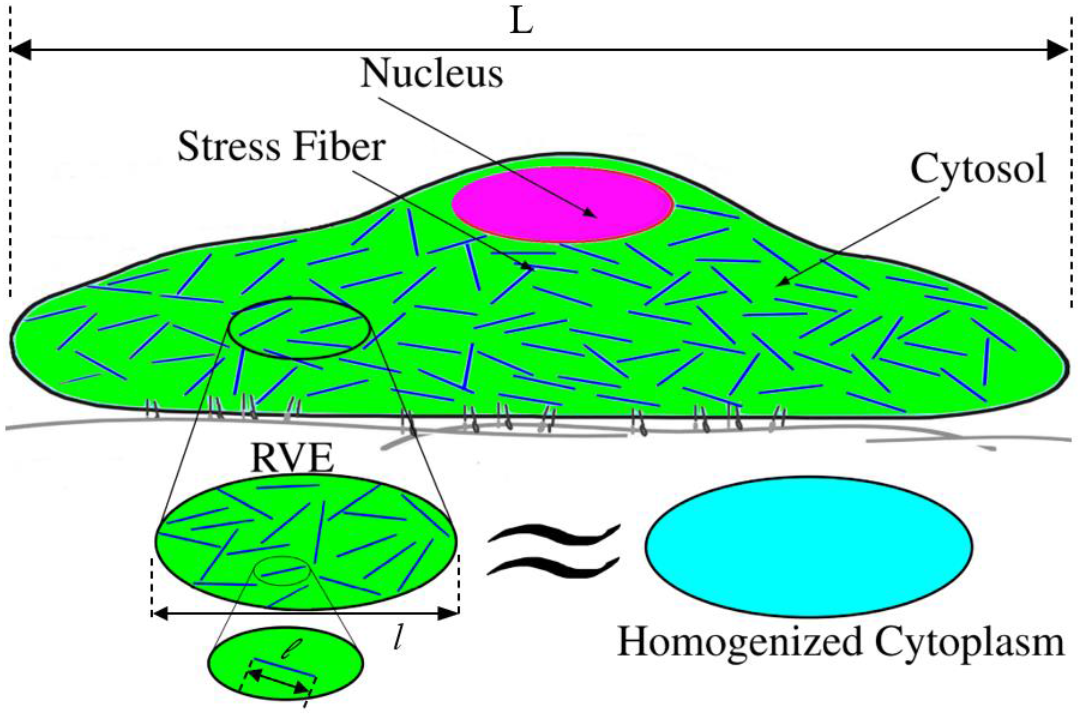
Schematic cross-section of the cell illustrating the random distribution of several actin stress fibres and the corresponding homogenized microstructure.

In microstructurally inhomogeneous materials, volume average stress and strain is obtained by integration of stress and strain over the RVE volume with respect to the microscopic coordinates inside the RVE:

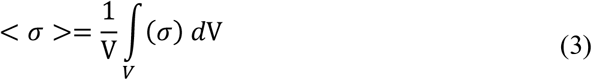

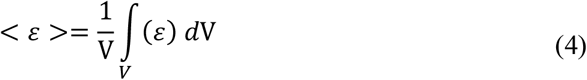

Here, σ(x) and ε(x) are the microscopic stress and strain, respectively, that are related to the average stress and strain by:

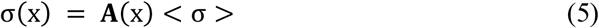

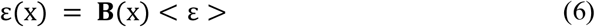

where **A** and **B** are the stress and strain concentration tensors, respectively.

Depending on the complexity of the microstructure, the concentration tensors may be obtained by different approximation approaches. The Mori-Tanaka homogenization scheme was used due to its superior accuracy for mid-range fibre volume fractions compared to the self-consistent approach, which is more accurate for high fibre volume fractions [59, 60]. A single ellipsoidal inclusion bonded to an infinite homogeneous elastic matrix subjected to uniform strain and stress at infinity was considered. For this problem, a suitable approximation approach is the mean field method, which is generally based on the Eshelby equivalent inclusion formulation [61]. The Mori-Tanaka (MT) homogenization model [62] is an effective field approximation based on Eshelby’s elasticity solution, assuming that the strain concentration tensor **B** is equal to the strain concentration of the single inclusion problem. The Mori-Tanaka method advances the Eshelby method, and the relationship for the effective strain is given as

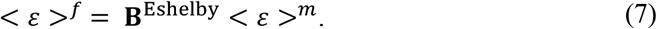

The concentration tensor **B**^Eshelby^ for Eshelby’s equivalent inclusion is

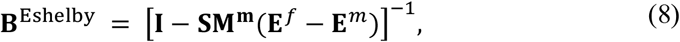

where **I** is the identity tensor, **S** is the Eshelby tensor [61], **E** is the elastic tensor, **M** is the compliance tensor, and *υ* is the volume fraction. The superscripts *f* and *m* refer to the fibre and matrix, respectively.

The Mori-Tanaka concentration tensor **B**^MT^ is given as

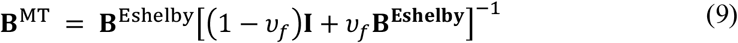

The relationship between the macroscopic stress < σ > and strain < ε > can be obtained by:

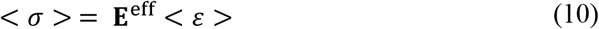

where **E**^eff^ is the effective elastic tensor of the homogeneous material obtained as a function of the strain concentration tensor **B**^MT^. The Neo-Hookean material model was employed to represent the homogenized cytoplasm’s non-linear stress-strain behaviour for large deformation.

The cytoplasm was considered as a composite of randomly oriented stress fibres in the cytosol representing the matrix. The cytosol was assumed hyperelastic with an elastic modulus of 1 kPa, see **Table 2**. The mechanical properties of stress fibres obtained from stretch tests [31] were utilized. The effective linear-elastic bulk modulus *K^eff^* and shear modulus *μ^eff^* of the cytosol matrix with randomly oriented and distributed stress fibres are given as:

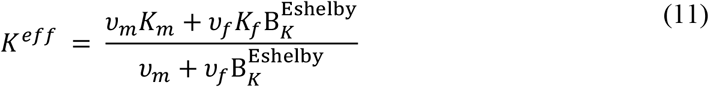

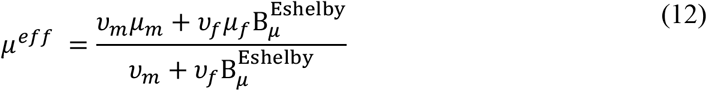

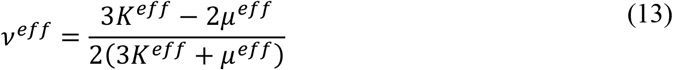

where *μ, K, ν*, and *υ* are the shear modulus, bulk modulus, Poisson’s ratio and volume fraction, respectively, of the fibre and matrix defined by subscripts *f* and *m*. The stress is obtained by:

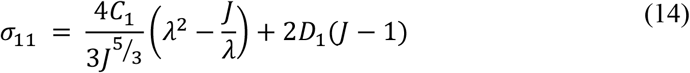

where C_1_ = *μ*/2, D_1_ = *K*/2, and *λ* is the principal stretch.

### 2.4 Parametric simulations

Changes in the actin cytoskeletal structure have been reported to affect cell fate [19]. The mechanical properties of the cellular components, including the nucleus, cytoplasm, actin cortex, and actin stress fibres, contribute to the cell’s effective mechanical stiffness [36]. Parametric simulations were conducted to determine the effect of stress fibre content and substrate elasticity on the cell’s deformation. The volume fraction of filamentous actin (F-actin) in the cytoskeleton has been reported to be approximately 1% [63]. As higher stress fibre volume fractions are expected in actin-rich intracellular regions (such as the cortex or actin bundles), stress fibre volume fraction of *υ*_f_ = 0, 1, 10, and 20% were used for the parametric simulations. **Figure 4** shows the hierarchical relation between micromechanical homogenization and implementation into finite element model.

**Figure 3.**
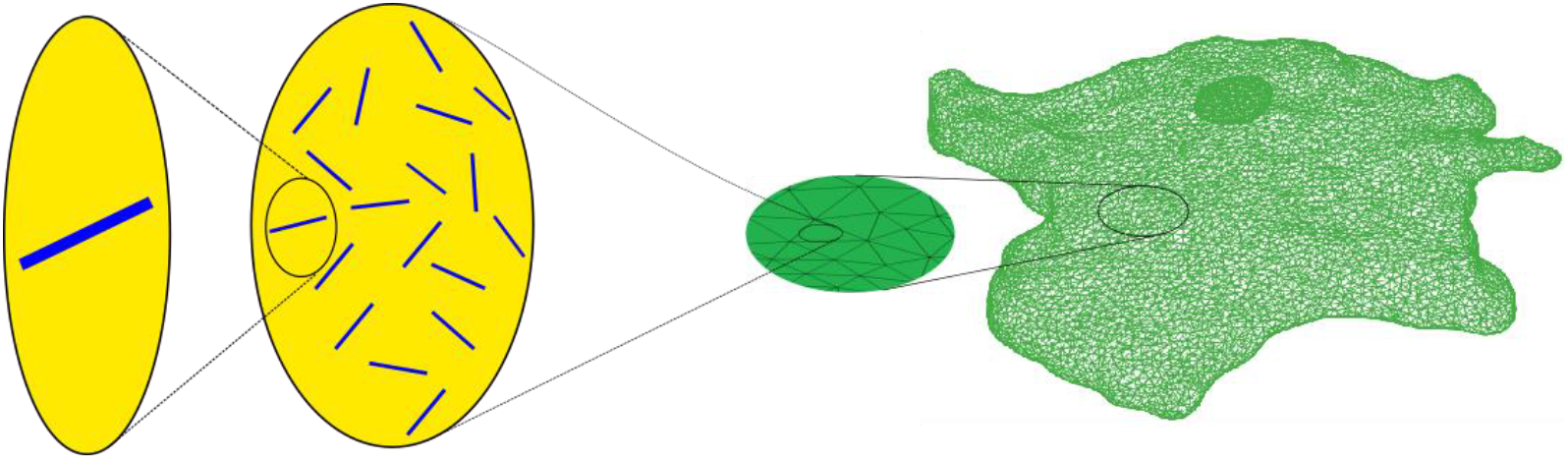
Schematic illustrating the upscaling from the Mori-Tanaka representative volume element to the whole-cell geometry of the finite element model.

For the substrate’s elastic modulus, the values of E_s_ = 0.01, 0.14, 1 and 10 MPa were used. The cell’s reference mechanical properties are summarised in **Table 2**.

### 2.5 Data analysis

The computed strain in the homogenized cytoplasm, nucleus and membrane of the cell exposed to uniaxial substrate stretch were recorded and assessed. Data are presented either for the mid-plane or the entire volume of the cell, respectively.

## 3. Results

### 3.1 Micromechanical homogenization of cytoplasm

The effective elastic modulus *E^eff^*, shear modulus *μ^eff^*, and bulk modulus *K^eff^* of the homogenized cytoplasm were predicted to increase, whereas the effective Poisson’s ratio *ν^eff^* decreased with increasing stress fibre volume fraction *υ*^f^, as illustrated in **Figure 4**.

**Figure 4.**
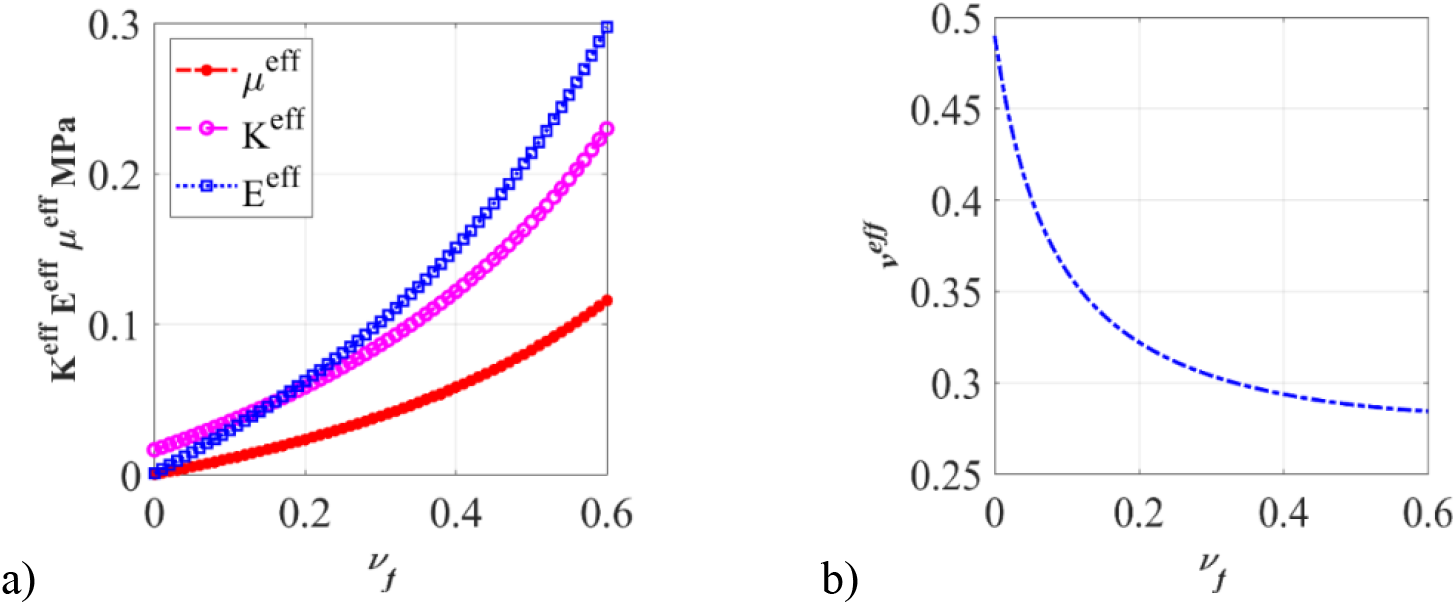
The effect of stress fibre volume fraction ν_f_ on (a) the effective elastic modulus **E**^eff^, shear modulus **μ**^eff^ and bulk modulus **K**^eff^, and (b) the effective Poisson’s ratio of the homogenized cytoplasm.

An increase in stress fibre volume fraction also resulted in an increase in the stress for a given stretch and the non-linearity of the stress-stretch relationship of the homogenized cytosol (**Figure 5**). The stress-stretch relationship can, however, be approximated as linear for small stretch values of λ ≤ 1.1.

**Figure 5.**
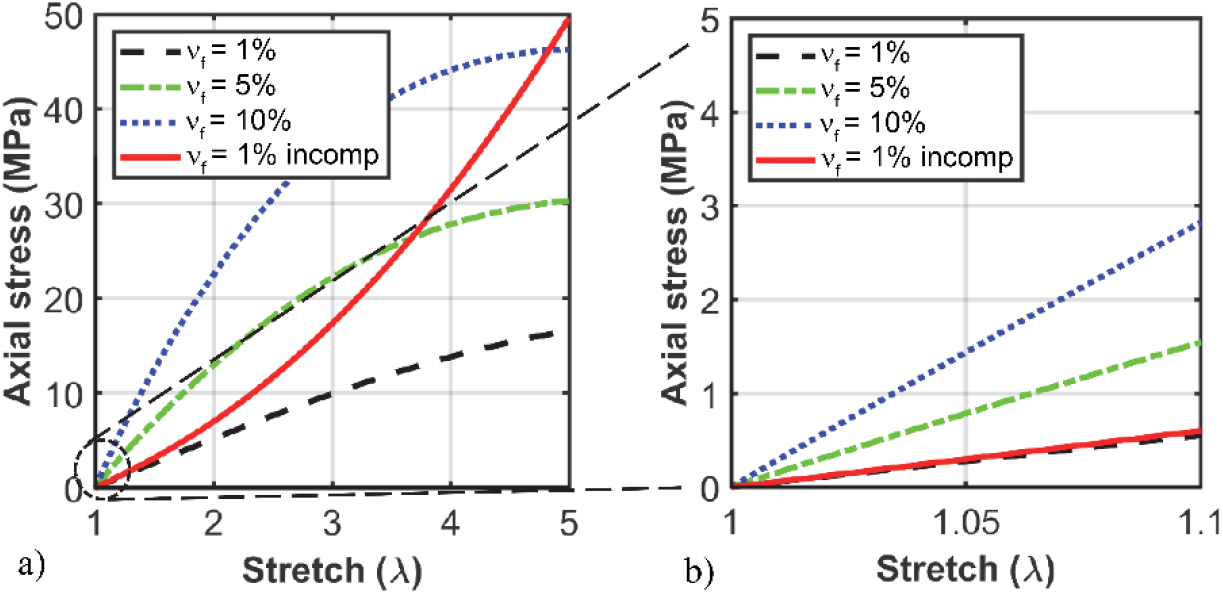
The axial stress as versus uniaxial stretch in the homogenized cytoplasm predicted with the hyperelastic Neo-Hookean material model for stress fibre volume fractions of ν_f_ = 1%, 5% and 10% and an incompressible case with ν_f_ = 1% for (a) large stretch λ = 1.0 to 5.0 and (b) small stretch λ = 1.0 to 1.1. (‘incomp’ refers to incompressible).

### 3.2 Effect of stress fibre volume fraction, substrate elastic modulus, and substrate deformation on intracellular deformation

The spatial distribution of the strain in the cytosol and nucleus was insensitive to the variation of the stress fibre volume fraction, whereas the strain magnitude decreased as the stress fibre content increased (**Figure 6**). The mid-plane peak strain decreased considerably with increasing stress fibre volume fraction in the cytoplasm and nucleus but only marginally in the cell membrane (**Figure 7**).

**Figure 6.**
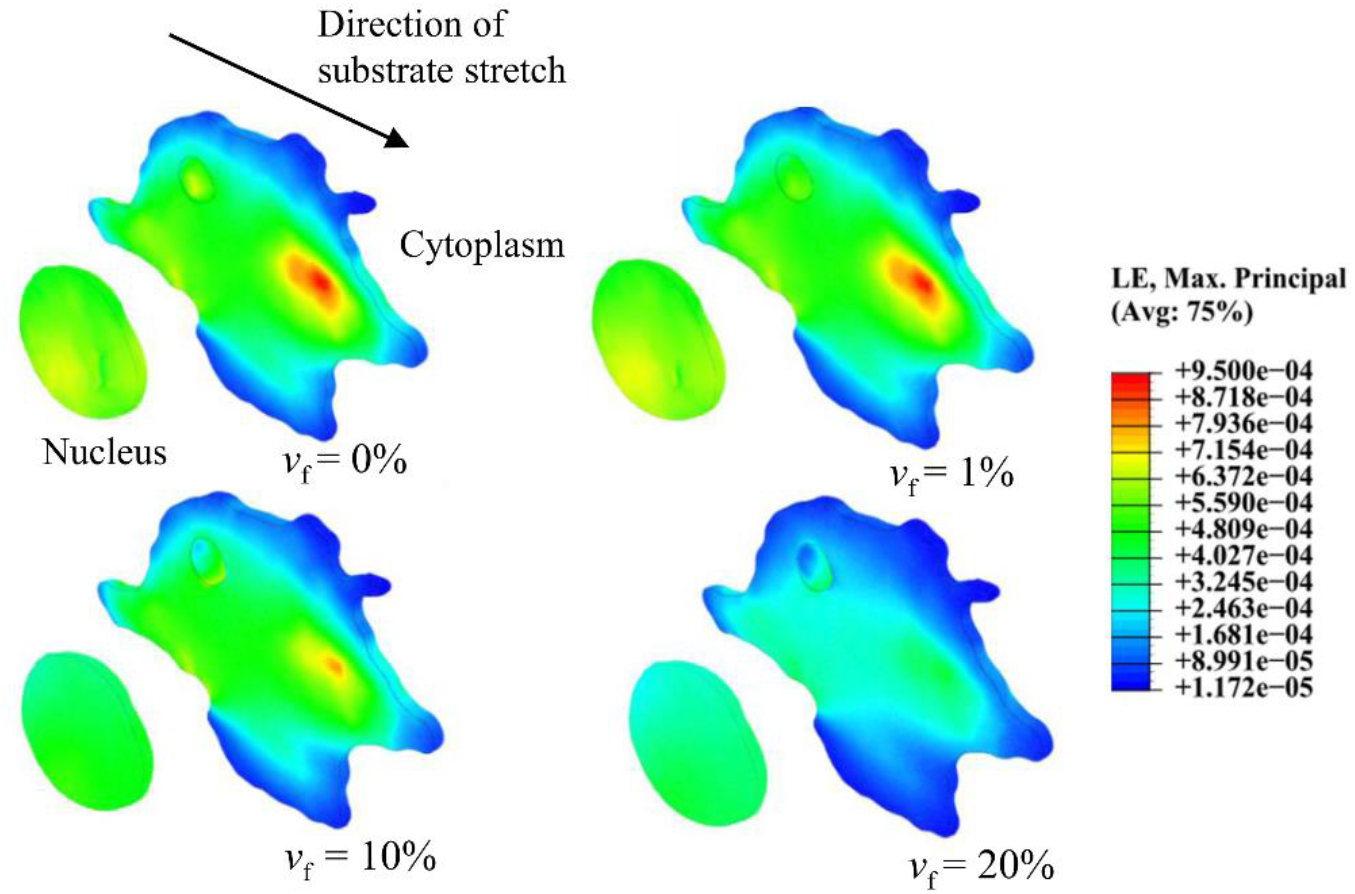
Spatial distribution and magnitude of maximum principal strain in the mid-plane of cytoplasm and nucleus for different stress fibre volume fractions υ_f_ = 0, 1, 10 and 20% and elastic modulus of the substrate of E_sub_ = 0.01 MPa. The intracellular strain’s spatial distribution was insensitive to changes in stress fibre content, whereas the strain magnitude decreased with increasing stress fibre content. The results are presented in the cell’s mid-plane to disregard potential numerical localization effects of the contact between the cell and the substrate.

**Figure 7.**
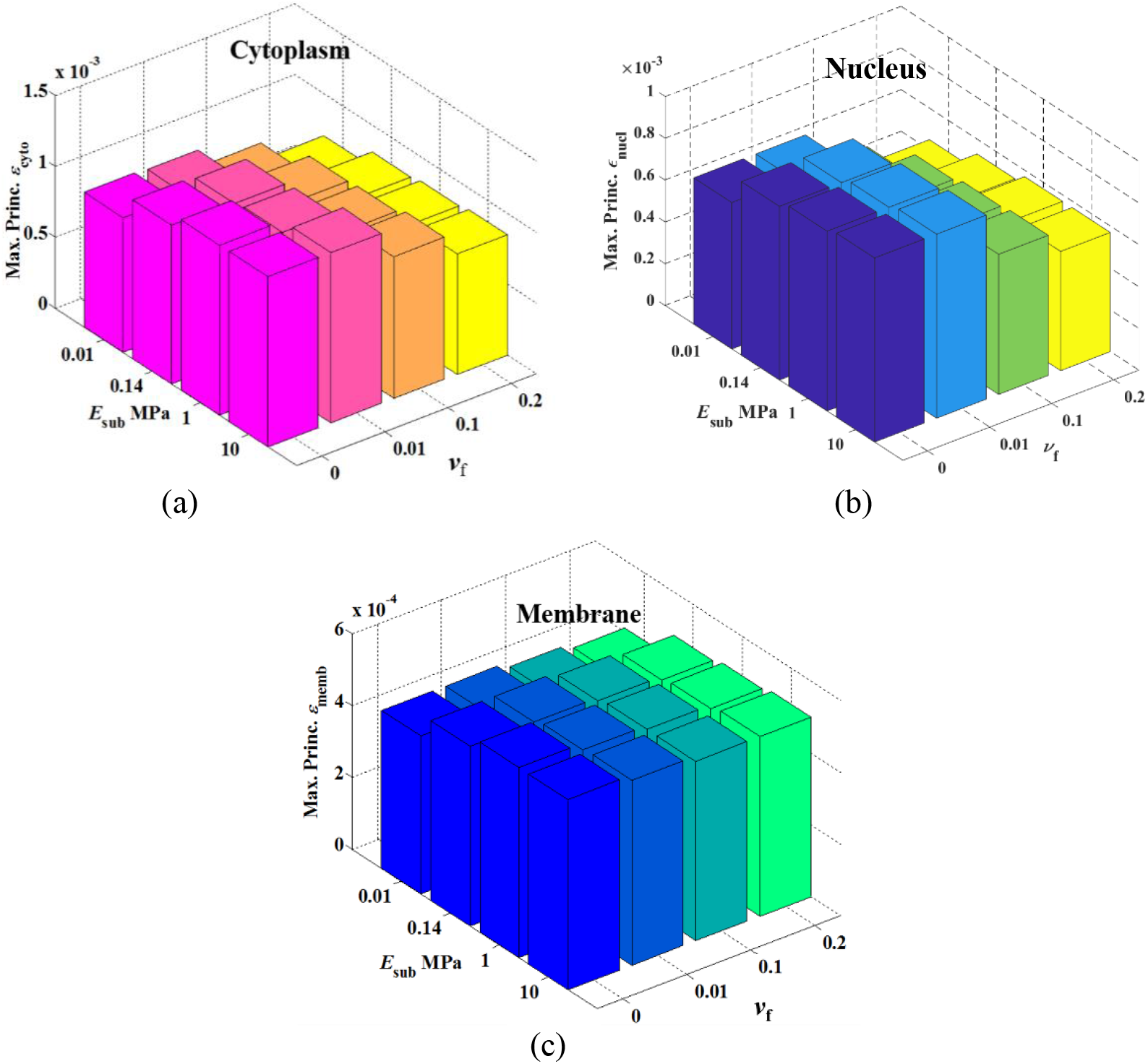
Peak maximum principal strain in the mid-section of the cytoplasm (a), nucleus (b) and membrane (c) at a substrate stretch of λ = 1.1 for different values of stress fibre volume fraction ν_f_ and substrate elastic modulus E_sub_. (The values of the peak maximum principal strains in the cytoplasm, nucleus and cell membrane are provided in the online supplement, **Table S1**).

An increase in the substrate’s elastic modulus E_sub_ led to an increase in the mid-plane peak strain in the nucleus, cytoplasm and membrane (**Figure 7**). The peak strain in the cellular components changed more for the variation of E_sub_ in the lower range (i.e. 0.01 and 0.14 MPa) compared to the higher values of E_sub_ (1 and 10 MPa). The lower values of E_sub_ are of the same order of magnitude as the elastic modulus of the cell components, whereas the higher values of E_sub_ considerably exceed those of the cell components.

The peak strain in the entire cytoplasm and nucleus, respectively, increased near-linearly with increasing peak strain in the substrate (**Figure 8** and **Figure 9**). (Note that strain in the entire cellular components is reported here compared to strain in the mid-plane section reported above.) The peak strains in cytoplasm and nucleus can be approximated from the substrate peak strain using the following linear functions and parameter values reported in **Table 3**:

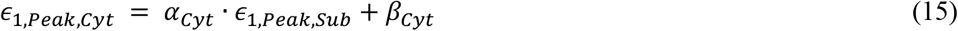

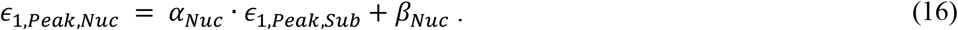

**Figure 8.**
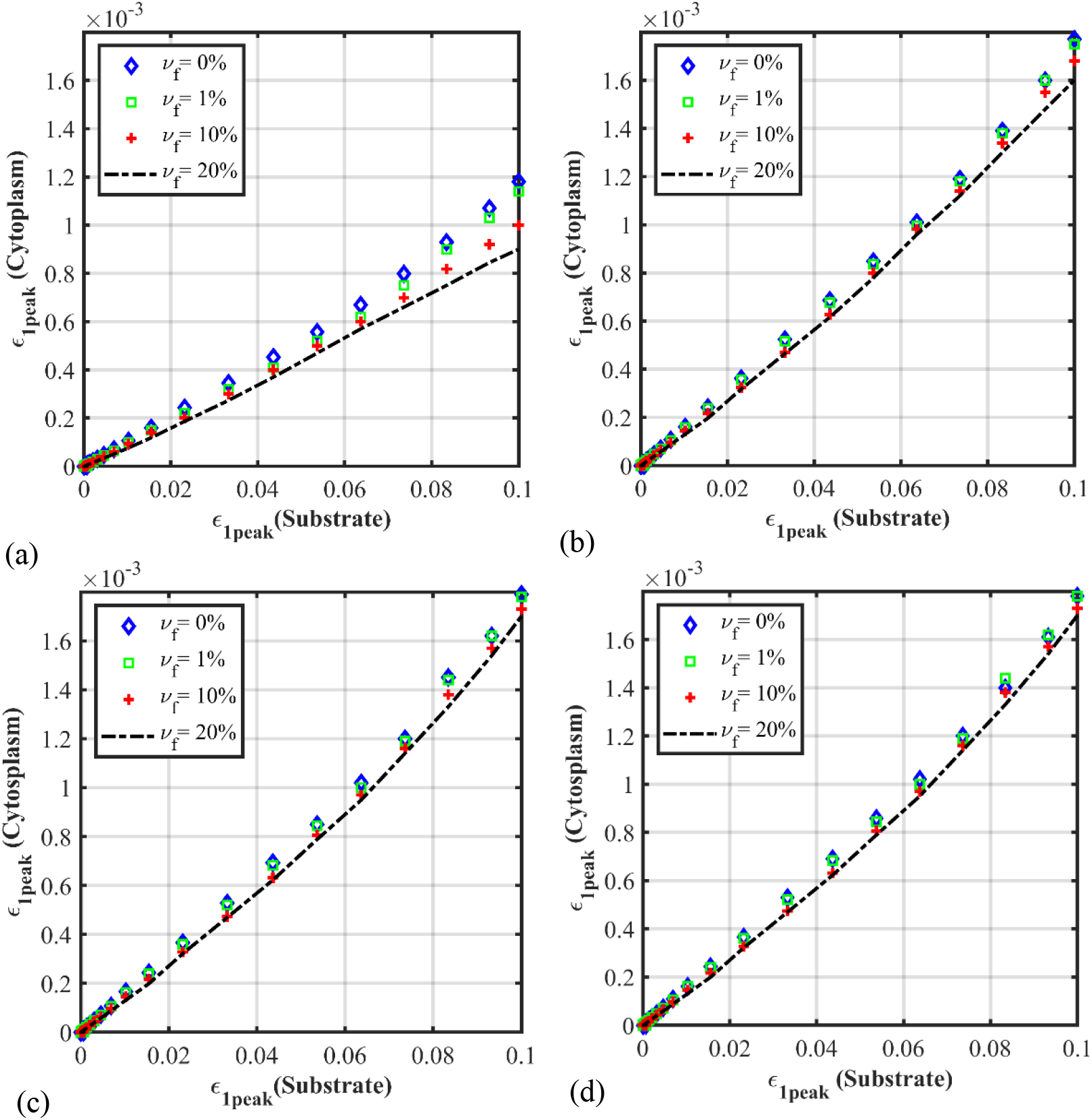
Peak maximum principal strain in the entire cytoplasm versus the substrate for different stress fibre volume fractions of 0, 1, 10 and 20% and substrate elastic modulus E_sub_ of 0.01 (a), 0.14 (b), 1 (c) and 10 MPa (d).

**Figure 9.**
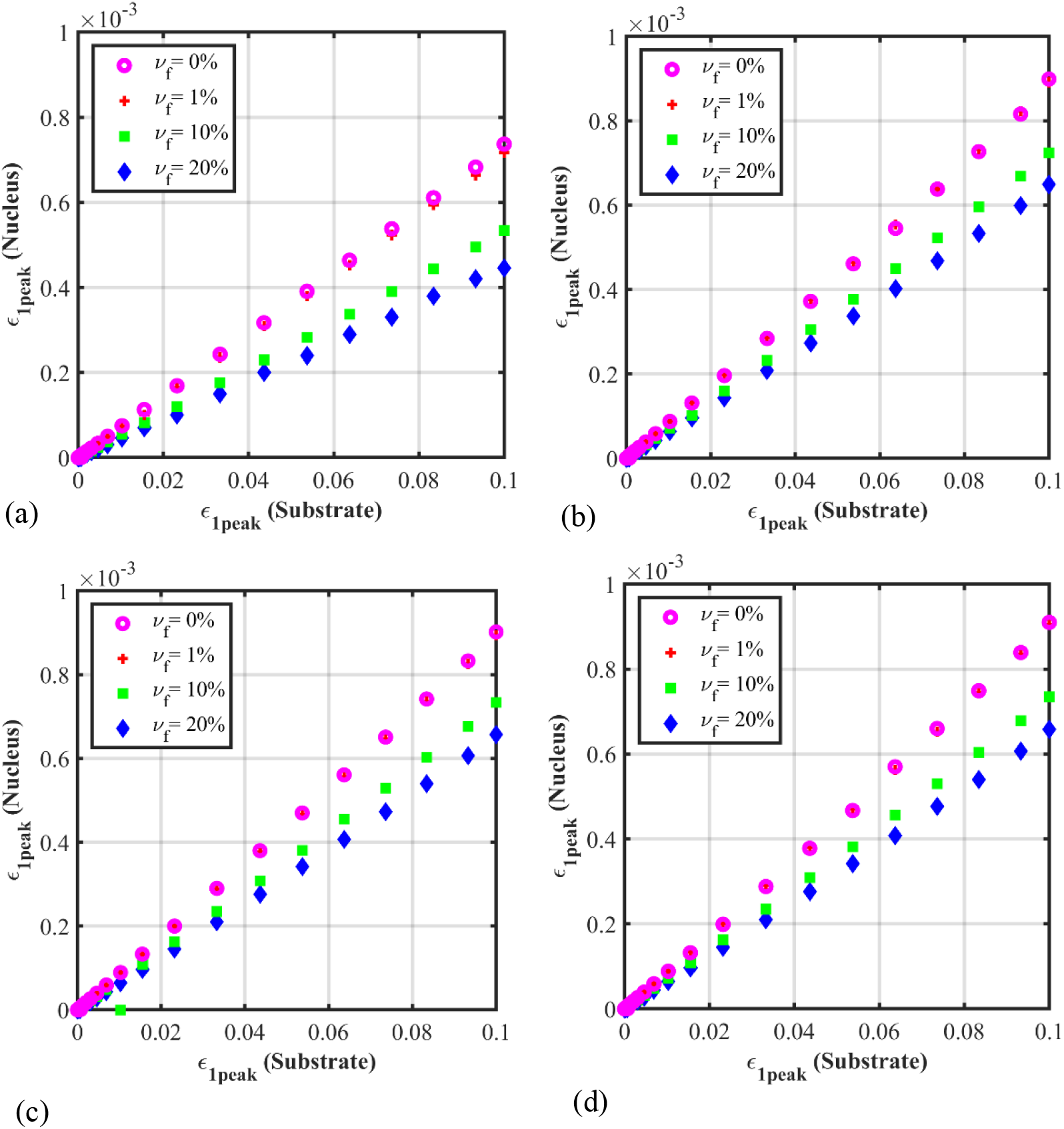
Peak maximum principal strain in the entire nucleus versus the substrate for different stress fibre volume fractions of 0, 1, 10 and 20% and substrate elastic modulus E_sub_ of 0.01 (a), 0.14 (b), 1 (c) and 10 MPa (d).

**Table 3.**
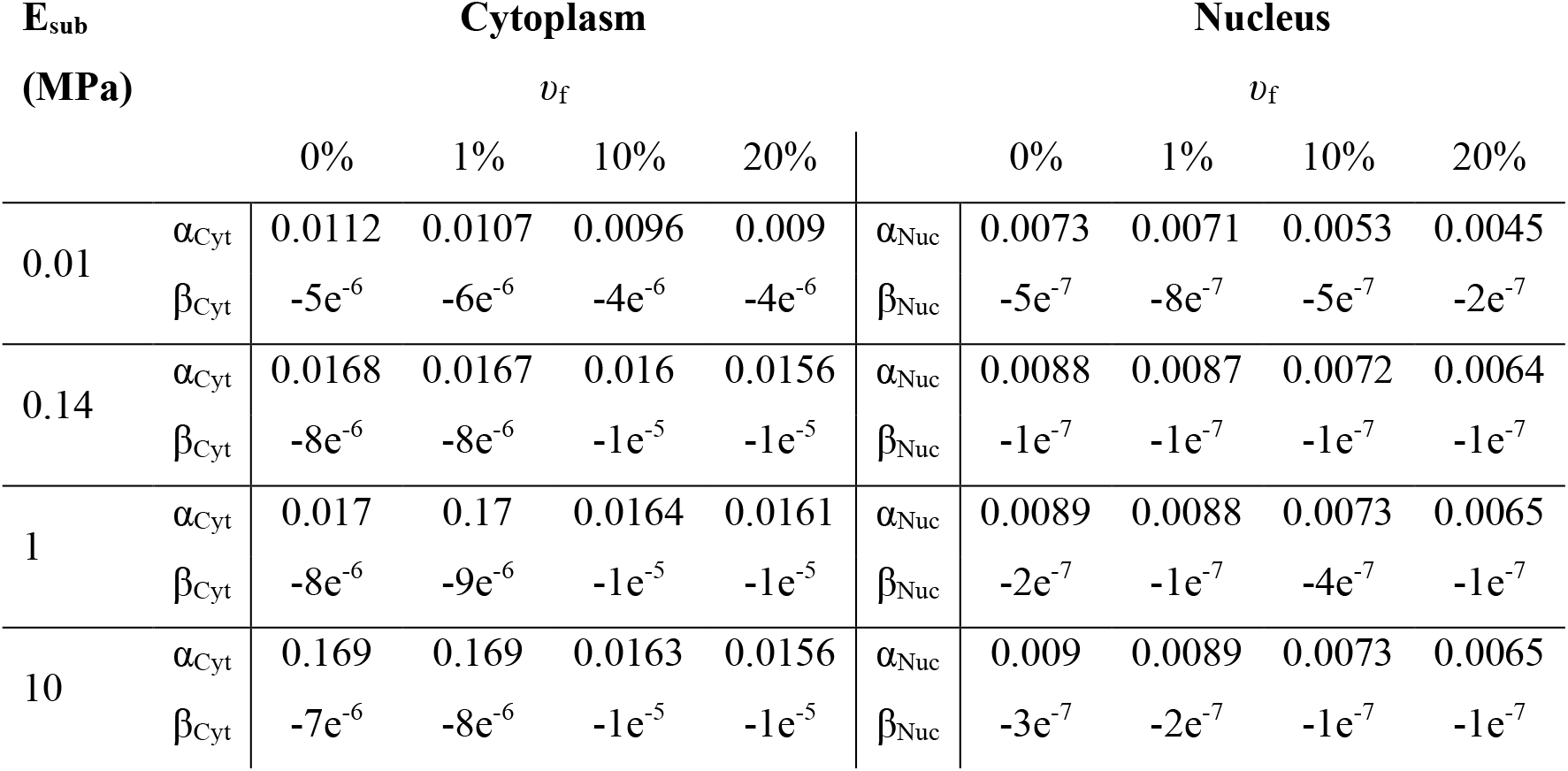
Parameter values to quantitatively predict peak strain in cytoplasm and nucleus (see Eqs. 15 and 16).

## 4. Discussion

Previous studies of eukaryotic cells have shown that the cytoskeletal structures largely determine the cytoplasm’s distinct mechanical properties. The assumption of homogenous mechanical properties for the entire cell has been a simplification in some computational studies of single-cell mechanics. This is particularly true for cells with focal adhesions at which stress fibres present critical inhomogeneities. Hybrid computational cell models [33–35] with a restricted number of tensegrity elements available to represent the mechanical contribution of stress fibres have remained limited in their capabilities to capture actual cellular behaviour. One of the main shortcomings of continuum-based models is the limited representation of the cytoskeletal fibres’ functional contribution [64].

In the current study, a finite element method and model for single-cell mechanics were developed that included the micromechanical homogenization of actin stress fibres as primary cytoskeletal elements in the cytoplasm. Using the model, the influence of stress fibre content on the deformation of cytoplasm, nucleus and cell membrane of a fibroblast was investigated for exposure to a uniaxial strain of up to 10% of substrates of different stiffness.

An increase in stress fibre volume fraction led to a decrease in the peak mid-plane strain in the cytoplasm and nucleus but affected the cell membrane’s strain to a lesser degree (**Figure 7**). The spatial strain distribution in the cytoplasm and nucleus was not affected by the change in stress fibre volume fraction, supposedly since the focal adhesions’ distribution was the same for all stress fibre volume fractions. We have investigated the effect of focal adhesions on intracellular strain distribution in previous studies [28, 65, 66].

The peak mid-plane strain in the cytoplasm and nucleus changed more for a change of the stress fibre volume fraction from 0% to 10% than from 10% to 20%. These results support the cytoskeleton’s significant role in transmitting extracellular mechanical forces to the nucleus, possibly mediating mechanotransduction [67]. The negligible change of the strain in the cell membrane indicates that the membrane is less sensitive to a change in stress fibre content than the cytosol and nucleus.

This study indicates that the omission of stress fibres in computational models may overestimate intracellular strain. Many computational studies used beam elements to represent stress fibres [8, 9, 11, 33] due to the challenges in discretely reconstructing the stress fibres. However, this is insufficient to capture the global effect of stress fibres for high volume fractions [68] and random orientations [69]. The approach utilizing micromechanical homogenization can address this current limitation in single-cell computational mechanics.

A substantial variation of the effective Poisson’s ratio of the micromechanically homogenized cytoplasm was predicted with the change in stress fibre content (**Figure 4**). This finding may support the wide range of values for Poisson’s ratio reported by previous studies, from nearly incompressible hyperelastic with 0.49 [70] to 0.3 [71, 72].

The model revealed that the maximum principal strain in the cytoplasm and nucleus is affected jointly by stress fibre content and substrate elastic modulus (Figure 4 and Figure 4). The predicted strains in the cytoplasm and nucleus agree reasonably with experimental data [73].

Once validated, the numerical relationships between peak strain in the cytoplasm and nucleus and the substrate strain (Eqs. 15 and 16) can facilitate the simple estimation of the maximum intracellular deformation for a given environmental stiffness, e.g. for guidance in engineering cellular microenvironments for therapeutic applications.

The cytoplasm’s hyperelastic response was captured by a Neo-Hookean strain energy density function based on isotropic elastic shear modulus and bulk modulus derived by micromechanical homogenization. Based on the fibroblast’s spread morphology with many stress fibre orientations observed microscopically, the homogenized cytoplasm was treated as mechanically isotropic with randomly aligned stress fibres.

There are some simplifications in the finite element model presented here. Focal adhesions were assumed to be arbitrarily distributed over the entire basal cell surface. We investigated the impact of focal adhesions in more detail in previous studies [28, 60, 61]. The cell-specific morphology of the model was intended to resemble only in part the in vivo interactions of the cell and substrate. This limitation and the model complexity were deemed acceptable for comparison with experimental studies of cellular stretching in adherent monolayers [74, 75, 76] and appropriate for the current study’s primary goal, namely to investigate the role of stress fibres on cellular mechanics with changing substrate stiffness.

Limitations of the current study include the absence of quantitative information on the stress fibre distribution and the omission of microtubules and intermediate filaments. The latter was due to the technical challenge to identify these structures in the microscopic images. The lack of validation of the model and the numerical results is based on a scarcity of experimental data for cellular and subcellular structural and mechanical properties and localized intracellular deformation. The parametric study of the substrate elastic modulus ignored the cells’ capability to adapt their mechanical and biochemical properties to environmental stiffness. These complex adaptations exceeded the scope of the current study.

Beyond addressing the limitations mentioned above, the extension of this study will include computationally modelling cytoskeletal dynamics, non-linear constitutive behaviour and intracellular viscosity.

## 5. Conclusion

This study demonstrated the importance of representing cytoskeletal stress fibres in computational models for single-cell mechanics. The findings can contribute to more realistic and accurate computational models for cell mechanics to improve the understanding of mechanotransduction in living cells and to developing a simple numerical tool to aid in designing engineered extracellular environments for therapeutic applications.

## Author biographies

**Tamer Abdalrahman** received his PhD from the Politecnico di Torino, Italy, in 2012 and subsequently undertook a postdoctoral fellowship at Vienna University of Technology, Austria. In 2015, he joined the Division of Biomedical Engineering at the University of Cape Town as postdoctoral research fellow to contribute to the research in therapies for myocardial infarction. His research focussed on cellular and peri-cellular mechanics in stem cell therapies for myocardial infarcts. Currently, he is senior postdoctoral fellow in the Computational Mechanobiology Group at the Julius Wolff Institute for Biomechanics and Musculoskeletal Regeneration, Charité – Universitätsmedizin Berlin. His current research focuses on the modeling of sprouting angiogenesis induction by cell-cell mechanical interaction.

**Neil Davies** is an Associate Professor in the Cardiovascular Research Unit in the Department of Surgery at the University of Cape Town. His research focusses on biomaterials as therapeutics for cardiovascular pathologies and medical devices. He has a significant interest in cellular biomechanics as well. The work in cardiovascular diseases primarily examines the influence of hydrogels as stand-alone treatments or as therapeutic agent vehicles for myocardial infarctions. A strong recent emphasis has been the regulation of cellular and tissue interactions with hydrogels. His research group uses a range of cell biology techniques and animal models to assay the impact of these therapeutic strategies.

**Thomas Franz** is Professor of Biomedical Engineering in the Department of Human Biology at the University of Cape Town. His research focusses on biomechanics and mechanobiology from organ to sub-cellular scale with application to cardiovascular diseases, cancers, and infectious diseases. The work in cardiovascular diseases includes biomaterial and cell therapies for myocardial infarction and heart failure and endothelial cell mechanics in disease. The research in cellular and subcellular mechanobiology aims at improving the understanding and treatment of cancers and studying interactions of virions and parasites with human host cells involved in infectious diseases such as chikungunya, influenza and malaria. His research group uses numerical, computational and experimental methods to probe mechanical properties and behaviour of biological structures in health and disease and to predict effects of therapeutic interventions.

## Acknowledgements

The research reported in this publication was supported by the National Research Foundation of South Africa (UID 92531 and 93542) and the South African Medical Research Council (SIR 328148). Views and opinions expressed are not those of the NRF or MRC but the authors.

## Availability of data

Abaqus input files of the finite element models used in this study are available on ZivaHUB (http://doi.org/10.25375/uct.9782798).

## References

[1] Huang, H., Kamm, R. D., & Lee, R. T. (2004). Cell mechanics and mechanotransduction: pathways, probes, and physiology. American Journal of Physiology-Cell Physiology, 287(1), C1–C11.

[2] Brown, T. D. (2000). Techniques for mechanical stimulation of cells in vitro: a review. Journal of biomechanics, 33(1), 3–14.

[3] Wang, N., Naruse, K., Stamenović, D., Fredberg, J. J., Mijailovich, S. M., Tolić-Nørrelykke, I. M., … & Ingber, D. E. (2001). Mechanical behavior in living cells consistent with the tensegrity model. Proceedings of the National Academy of Sciences, 98(14), 7765–7770.

[4] Moeendarbary, E., Harris, A. R. (2014). Cell mechanics: principles, practices, and prospects. Wiley Interdisciplinary Reviews: Systems Biology and Medicine, 6(5), 371–388.

[5] Hu, S., Chen, J., Fabry, B., Numaguchi, Y., Gouldstone, A., Ingber, D. E., … & Wang, N. (2003). Intracellular stress tomography reveals stress focusing and structural anisotropy in cytoskeleton of living cells. American Journal of Physiology-Cell Physiology, 285(5), C1082–C1090.

[6] Karcher, H., Lammerding, J., Huang, H., Lee, R. T., Kamm, R. D., & Kaazempur-Mofrad, M. R. (2003). A three-dimensional viscoelastic model for cell deformation with experimental verification. Biophysical journal, 85(5), 3336–3349.

[7] Ohayon, J., Tracqui, P. (2005). Computation of adherent cell elasticity for critical cell-bead geometry in magnetic twisting experiments. Annals of biomedical engineering, 33(2), 131–141.

[8] Evans, N. D., Gentleman, E. (2014). The role of material structure and mechanical properties in cell–matrix interactions. Journal of Materials Chemistry B, 2(17), 2345–2356.

[9] Barreto, S., Clausen, C. H., Perrault, C. M., Fletcher, D. A., Lacroix, D. (2013). A multi-structural single cell model of force-induced interactions of cytoskeletal components. Biomaterials, 34(26), 6119–6126.

[10] Barreto, S., Perrault, C. M., Lacroix, D. (2014). Structural finite element analysis to explain cell mechanics variability. Journal of the Mechanical Behavior of biomedical materials, 38, 219–231.

[11] Ishiko, A., Shimizu, H., Kikuchi, A., Ebihara, T., Hashimoto, T., Nishikawa, T. (1993). Human autoantibodies against the 230-kD bullous pemphigoid antigen (BPAG1) bind only to the intracellular domain of the hemidesmosome, whereas those against the 180-kD bullous pemphigoid antigen (BPAG2) bind along the plasma membrane of the hemidesmosome in normal human and swine skin. The Journal of clinical investigation, 91(4), 1608–1615.

[12] homas, G., Burnham, N. A., Camesano, T. A., & Wen, Q. (2013). Measuring the mechanical properties of living cells using atomic force microscopy. Journal of Visualized Experiments 27(76): e50497.

[13] Park, C. Y., Tambe, D., Alencar, A. M., Trepat, X., Zhou, E. H., Millet, E., Butler, J.P., Fredberg, J. J. (2010). Mapping the cytoskeletal prestress. American Journal of Physiology-Cell Physiology, 298(5), C1245–C1252.

[14] Sosa, M. S., Bragado, P., Aguirre-Ghiso, J. A. (2014). Mechanisms of disseminated cancer cell dormancy: an awakening field. Nature Reviews Cancer, 14(9), 611–622.

[15] Mavrakis, M., Azou-Gros, Y., Tsai, F. C., Alvarado, J., Bertin, A., Iv, F., Kress, A., Brasselet, S., Koenderink, G.H., Lecuit, T. (2014). Septins promote F-actin ring formation by crosslinking actin filaments into curved bundles. Nature cell biology, 16(4), 322–334.

[16] Lancaster, O. M., Baum, B. (2014). Shaping up to divide: coordinating actin and microtubule cytoskeletal remodelling during mitosis. In Seminars in cell & developmental biology 34, 109–115.

[17] Stewart, M. P., Toyoda, Y., Hyman, A. A., & Müller, D. J. (2012). Tracking mechanics and volume of globular cells with atomic force microscopy using a constant-height clamp. nature protocols, 7(1), 143–154.

[18] Rotsch, C., Radmacher, M. (2000). Drug-induced changes of cytoskeletal structure and mechanics in fibroblasts: an atomic force microscopy study. Biophysical journal, 78(1), 520–535.

[19] Fletcher, D. A., Mullins, R. D. (2010). Cell mechanics and the cytoskeleton. Nature, 463(7280), 485–492.

[20] Wu, H. W., Kuhn, T., Moy, V. T. (1998). Mechanical properties of L929 cells measured by atomic force microscopy: effects of anticytoskeletal drugs and membrane crosslinking. Scanning: The Journal of Scanning Microscopies, 20(5), 389–397.

[21] Janmey, P. A., Miller, R. T. (2011). Mechanisms of mechanical signaling in development and disease. Journal of cell science, 124(1), 9–18.

[22] Lekka, M., Laidler, P., Gil, D., Lekki, J., Stachura, Z., Hrynkiewicz, A. Z. (1999). Elasticity of normal and cancerous human bladder cells studied by scanning force microscopy. European Biophysics Journal, 28(4), 312–316.

[23] Na, S., Sun, Z., Meininger, G. A., Humphrey, J. D. (2004). On atomic force microscopy and the constitutive behavior of living cells. Biomechanics and Modeling in Mechanobiology, 3(2), 75–84.

[24] Humphrey, J. D. (2002). On mechanical modeling of dynamic changes in the structure and properties of adherent cells. Mathematics and Mechanics of Solids, 7(5), 521–539.

[25] Lim, C. T., Zhou, E. H., Quek, S. T. (2006). Mechanical models for living cells-a review. Journal of biomechanics, 39(2), 195–216.

[26] Lavagnino, M., Arnoczky, S. P., Kepich, E., Caballero, O., Haut, R. C. (2008). A finite element model predicts the mechanotransduction response of tendon cells to cyclic tensile loading. Biomechanics and modeling in mechanobiology, 7(5), 405–416.

[27] Miller, P., Hu, L., Wang, J. (2010). Finite element simulation of cell–substrate decohesion by laser-induced stress waves. Journal of the mechanical behavior of biomedical materials, 3(3), 268–277.

[28] Abdalrahman, T., Dubuis, L., Green, J., Davies, N., Franz, T. (2017). Cellular mechanosensitivity to substrate stiffness decreases with increasing dissimilarity to cell stiffness. Biomechanics and modeling in mechanobiology, 16(6), 2063–2075.

[29] Simon, B. R., Kaufmann, M. V., McAfee, M. A., Baldwin, A. L. (1993). Finite element models for arterial wall mechanics. Journal of Biomechanical Engineering 115:489–496.

[30] Milner, J. S., Grol, M. W., Beaucage, K. L., Dixon, S. J., Holdsworth, D. W. (2012). Finite-element modeling of viscoelastic cells during high-frequency cyclic strain. Journal of functional biomaterials, 3(1), 209–224.

[31] Mullen, C. A., Vaughan, T. J., Voisin, M. C., Brennan, M. A., Layrolle, P., McNamara, L. M. (2014). Cell morphology and focal adhesion location alters internal cell stress. Journal of The Royal Society Interface, 11(101), 20140885.

[32] Verbruggen, S. W., Vaughan, T. J., McNamara, L. M. (2012). Strain amplification in bone mechanobiology: a computational investigation of the in vivo mechanics of osteocytes. Journal of the Royal Society Interface, 9(75), 2735–2744.

[33] Slomka, N., Gefen, A. (2010). Confocal microscopy-based three-dimensional cell-specific modeling for large deformation analyses in cellular mechanics. Journal of biomechanics, 43(9), 1806–1816.

[34] Slomka, N., Gefen, A. (2012). Relationship between strain levels and permeability of the plasma membrane in statically stretched myoblasts. Annals of biomedical engineering, 40(3), 606–618.

[35] Yao, Y., Lacroix, D., Mak, A. F. (2016). Effects of oxidative stress-induced changes in the actin cytoskeletal structure on myoblast damage under compressive stress: confocal-based cell-specific finite element analysis. Biomechanics and modeling in mechanobiology, 15(6), 1495–1508.

[36] Chen, A., Moy, V. T. (2000). Cross-linking of cell surface receptors enhances cooperativity of molecular adhesion. Biophysical Journal, 78(6), 2814–2820.

[37] Hine R (2009) The Facts on File Dictionary of Biology. Infobase Publishing.

[38] Fortino, S., Zagari, G., Mendicino, A. L., Dill-Langer, G. (2012). A simple approach for FEM simulation of Mode I cohesive crack growth in glued laminated timber under short-term loading. Journal of Structural Mechanics, 45(1), 1–20.

[39] Bartalena, G., Loosli, Y., Zambelli, T., Snedeker, J. G. (2012). Biomaterial surface modifications can dominate cell–substrate mechanics: the impact of PDMS plasma treatment on a quantitative assay of cell stiffness. Soft Matter, 8(3), 673–681.

[40] Balaban, N.Q., Schwarz, U.S., Riveline, D., Goichberg, P., Tzur, G., Sabanay, I., Mahalu, D., Safran, S., Bershadsky, A., Addadi, L. and Geiger, B. (2001). Force and focal adhesion assembly: a close relationship studied using elastic micropatterned substrates. Nature cell biology, 3(5), 466–472.

[41] Goffin, J. M., Pittet, P., Csucs, G., Lussi, J. W., Meister, J. J., Hinz, B. (2006). Focal adhesion size controls tension-dependent recruitment of α-smooth muscle actin to stress fibers. The Journal of cell biology, 172(2), 259–268.

[42] Ishiko, A., Shimizu, H., Kikuchi, A., Ebihara, T., Hashimoto, T., Nishikawa, T. (1993). Human autoantibodies against the 230-kD bullous pemphigoid antigen (BPAG1) bind only to the intracellular domain of the hemidesmosome, whereas those against the 180-kD bullous pemphigoid antigen (BPAG2) bind along the plasma membrane of the hemidesmosome in normal human and swine skin. The Journal of clinical investigation, 91(4), 1608–1615.

[43] Park, K., Paulino, G. H. (2012). Computational implementation of the PPR potential-based cohesive model in ABAQUS: Educational perspective. Engineering fracture mechanics, 93, 239–262.

[44] Dávila, C. G., Camanho, P. P., Turon, A. (2008). Effective simulation of delamination in aeronautical structures using shells and cohesive elements. Journal of Aircraft, 45(2), 663–672.

[45] Engler, A. J., Griffin, M. A., Sen, S., Bonnemann, C. G., Sweeney, H. L., Discher, D. E. (2004). Myotubes differentiate optimally on substrates with tissue-like stiffness pathological implications for soft or stiff microenvironments. Journal of Cell Biology, 166(6), 877–887.

[46] Breuls, R. G., Sengers, B. G., Oomens, C. W., Bouten, C. V., Baaijens, F. P. (2002). Predicting local cell deformations in engineered tissue constructs: a multilevel finite element approach. J. Biomech. Eng., 124(2), 198–207.

[47] Baaijens, F. P., Trickey, W. R., Laursen, T. A., Guilak, F. (2005). Large deformation finite element analysis of micropipette aspiration to determine the mechanical properties of the chondrocyte. Annals of biomedical engineering, 33(4), 494–501.

[48] Peeters, E. A. G., Oomens, C. W. J., Bouten, C. V. C., Bader, D. L., Baaijens, F. P. T. (2005). Mechanical and failure properties of single attached cells under compression. Journal of biomechanics, 38(8), 1685–1693.

[49] Jean, R. P., Chen, C. S., Spector, A. A. (2005). Finite-element analysis of the adhesion-cytoskeleton-nucleus mechanotransduction pathway during endothelial cell rounding: axisymmetric model. Journal of Biomechanical Engineering, 127(4), 594–600.

[50] Hochmuth, R. M., Mohandas, N., Blackshear Jr, P. L. (1973). Measurement of the elastic modulus for red cell membrane using a fluid mechanical technique. Biophysical journal, 13(8), 747–762.

[51] Lelidis, I., Joanny, J. F. (2013). Interaction of focal adhesions mediated by the substrate elasticity. Soft Matter, 9(46), 11120–11128.

[52] Nicolas, A., Safran, S. A. (2006). Limitation of cell adhesion by the elasticity of the extracellular matrix. Biophysical journal, 91(1), 61–73.

[53] Maniotis, A. J., Chen, C. S., & Ingber, D. E. (1997). Demonstration of mechanical connections between integrins, cytoskeletal filaments, and nucleoplasm that stabilize nuclear structure. Proceedings of the National Academy of Sciences, 94(3), 849–854.

[54] Wang, N., & Stamenovic, D. (2000). Contribution of intermediate filaments to cell stiffness, stiffening, and growth. American Journal of Physiology-Cell Physiology, 279(1), C188–C194.

[55] Brangwynne, C. P., MacKintosh, F. C., Kumar, S., Geisse, N. A., Talbot, J., Mahadevan, L., … & Weitz, D. A. (2006). Microtubules can bear enhanced compressive loads in living cells because of lateral reinforcement. The Journal of cell biology, 173(5), 733–741.

[56] Alberts, B., Johnson, A., Lewis, J., Raff, M., Roberts, K., & Walter, P. (2002). Integrins. In Molecular Biology of the Cell. 4th edition. Garland Science.

[57] Heidemann, S. R., Wirtz, D. (2004). Towards a regional approach to cell mechanics. Trends in cell biology, 14(4), 160–166.

[58] Feneberg, W., Aepfelbacher, M., & Sackmann, E. (2004). Microviscoelasticity of the apical cell surface of human umbilical vein endothelial cells (HUVEC) within confluent monolayers. Biophysical journal, 87(2), 1338–1350.

[59] Bakkali, A., Azrar, L., Fakri, N. (2011). Modeling of effective properties of multiphase magnetoelectroelastic heterogeneous materials. Computers, Materials, & Continua, 23(3), 201–231.

[60] Ryu, S., Lee, S., Jung, J., Lee, J., Kim, Y. (2019). Micromechanics-based homogenization of the effective physical properties of composites with an anisotropic matrix and interfacial imperfections. Frontiers in Materials, 6, 21.

[61] Eshelby, J. D. (1957). The determination of the elastic field of an ellipsoidal inclusion, and related problems. Proceedings of the royal society of London. Series A. Mathematical and physical sciences, 241(1226), 376–396.

[62] Mori, T., & Tanaka, K. (1973). Average stress in matrix and average elastic energy of materials with misfitting inclusions. Acta metallurgica, 21(5), 571–574.

[63] Satcher Jr, R. L., & Dewey Jr, C. F. (1996). Theoretical estimates of mechanical properties of the endothelial cell cytoskeleton. Biophysical journal, 71(1), 109–118.

[64] Ingber, D. E., Heidemann, S. R., Lamoureux, P., and Buxbaum, R. E., (2000). Opposing Views on Tensegrity as a Structural Framework for Understanding Cell Mechanics,” J. Appl. Physiol., 89, 1663–1678.

[65] Kohn, J. C., Abdalrahman, T., Sack, K. L., Reinhart-King, C. A., & Franz, T. (2018). Endothelial cells on an aged subendothelial matrix display heterogeneous strain profiles in silico. Biomechanics and modeling in mechanobiology, 17(5), 1405–1414.

[66] Kohn, J. C., Abdalrahman, T., Sack, K. L., Reinhart-King, C. A., & Franz, T. (2019). Cell focal adhesion clustering leads to decreased and homogenized basal strains. International journal for numerical methods in biomedical engineering, 35(12), e3260.

[67] Dahl, K. N., Ribeiro, A. J., & Lammerding, J. (2008). Nuclear shape, mechanics, and mechanotransduction. Circulation research, 102(11), 1307–1318.

[68] Vernerey, F. J., Farsad, M. (2014). A mathematical model of the coupled mechanisms of cell adhesion, contraction and spreading. Journal of mathematical biology, 68(4), 989–1022.

[69] Petroll, W. M., Cavanagh, H. D., Barry, P., Andrews, P., & Jester, J. V. (1993). Quantitative analysis of stress fiber orientation during corneal wound contraction. Journal of Cell Science, 104(2), 353–363.

[70] Sbrana, F., Sassoli, C., Meacci, E., Nosi, D., Squecco, R., Paternostro, F., … & Formigli, L. (2008). Role for stress fiber contraction in surface tension development and stretch-activated channel regulation in C2C12 myoblasts. American Journal of Physiology-Cell Physiology, 295(1), C160–C172.

[71] Caille, N., Thoumine, O., Tardy, Y., & Meister, J. J. (2002). Contribution of the nucleus to the mechanical properties of endothelial cells. Journal of biomechanics, 35(2), 177–187.

[72] Ferko, M. C., Bhatnagar, A., Garcia, M. B., & Butler, P. J. (2007). Finite-element stress analysis of a multicomponent model of sheared and focally-adhered endothelial cells. Annals of biomedical engineering, 35(2), 208–223.

[73] Gilchrist, C. L., Witvoet-Braam, S. W., Guilak, F., & Setton, L. A. (2007). Measurement of intracellular strain on deformable substrates with texture correlation. Journal of biomechanics, 40(4), 786–794.

[74] Schaffer, J. L., Rizen, M., L’Italien, G. J., Benbrahim, A., Megerman, J., Gerstenfeld, L. C., & Gray, M. L. (1994). Device for the application of a dynamic biaxially uniform and isotropic strain to a flexible cell culture membrane. Journal of Orthopaedic Research, 12(5), 709–719.

[75] Park, J. S., Chu, J. S., Cheng, C., Chen, F., Chen, D., & Li, S. (2004). Differential effects of equiaxial and uniaxial strain on mesenchymal stem cells. Biotechnology and bioengineering, 88(3), 359–368.

[76] Wang, N. (2010). Structural basis of stress concentration in the cytoskeleton. Molecular & Cellular Biomechanics, 7(1), 33–44.

