## Supplemental Table S1 for "In silico stress fibre content affects peak strain in cytoplasm and nucleus but not in the membrane for uniaxial substrate stretch"

### Online Supplement

**Table S1**. Peak maximum principal strain in the nucleus and cytoplasm at a substrate stretch of λ = 1.1 for different values of elastic modulus of substrate and stress fibre volume fraction.

| $f_{SF}$ | Peak Strain in Nucleus | | | | Peak Strain in Cytoplasm | | | |
|---|---|---|---|---|---|---|---|---|
| | $E_{sub}$ (MPa) | | | | $E_{sub}$ (MPa) | | | |
| | 0.01 | 0.14 | 1 | 10 | 0.01 | 0.14 | 1 | 10 |
| 0% | $7.37e^{-4}$ | $8.83e^{-4}$ | $9.02e^{-4}$ | $9.10e^{-4}$ | $1.18e^{-3}$ | $1.77e^{-3}$ | $1.79e^{-3}$ | $1.79e^{-3}$ |
| 1% | $7.17e^{-4}$ | $8.81e^{-4}$ | $8.94e^{-4}$ | $9.04e^{-4}$ | $1.14e^{-3}$ | $1.75e^{-3}$ | $1.78e^{-3}$ | $1.78e^{-3}$ |
| 10% | $5.34e^{-4}$ | $7.24e^{-4}$ | $7.34e^{-4}$ | $7.35e^{-4}$ | $1.00e^{-3}$ | $1.68e^{-3}$ | $1.73e^{-3}$ | $1.76e^{-3}$ |
| 20% | $4.46e^{-4}$ | $6.49e^{-4}$ | $6.57e^{-4}$ | $6.58e^{-4}$ | $0.90e^{-3}$ | $1.60e^{-3}$ | $1.70e^{-3}$ | $1.73e^{-3}$ |
